# Autochthonous faecal virome transplantation (FVT) reshapes the murine microbiome after antibiotic perturbation

**DOI:** 10.1101/591099

**Authors:** Lorraine A. Draper, Feargal J. Ryan, Marion Dalmasso, Pat G. Casey, Angela McCann, Vimalkumar Velayudhan, R. Paul Ross, Colin Hill

**Affiliations:** APC Microbiome Ireland, University College Cork, Cork, Ireland; School of Microbiology, University College Cork, Cork, Ireland; Teagasc Food Research Centre, Moorepark, Fermoy, Co. Cork, Ireland

**Keywords:** Bacteriophage, Virome, Transplantation, Microbiome, Antibiotic, Murine, Bacteriome

## Abstract

**Background:** It has become increasingly apparent that establishing and maintaining a complex and diverse gut microbiome is fundamental to human health. There are growing efforts to identify methods that can modulate and influence the microbiome, especially in individuals who due to disease or circumstance have experienced a disruption in their native microbiome. Faecal microbial transplantation (FMT) is one method that restores diversity to the microbiome of an individual by introducing microbes from a healthy donor. FMT introduces a complete microbiome into the recipient, including the bacteriome, archaeome, mycome and virome. In this study we investigated whether transplanting an autochthonous faecal virome consisting primarily of bacteriophages could impact a bacteriome disrupted by antibiotic treatment (**F**aecal **V**irome **T**ransplantation; **FVT**).

**Results:** Following disruption of the bacteriome by penicillin and streptomycin, test mice (n=8) received a bacteria free, faecal transplant, while Control mice (n=8) received a heated and nuclease treated control. The bacteriomes (as determined via 16S rRNA sequencing) of mice that received an FVT, in which bacteriophages predominate, separated from those of the Control mice as determined by principle co-ordinate analysis (PCoA), and contained differentially abundant taxa that reshaped the bacteriome profile such that it more closely resembled that of the pre-treatment mice. Similarly, metagenomic sequencing of the virome confirmed that the bacteriophages present in the gut of treatment and Control mice differed over time in both abundance and diversity, with transplanted phages seen to colonise the FVT mice.

**Conclusions:** An autochthonous virome transplant impacts on the bacteriome and virome of mice following antibiotic treatment. The virome, consisting mainly of bacteriophages, reshapes the bacteriome such that it more closely resembles the pre-antibiotic state. To date, faecal transplants have largely focussed on transferring living microbes, but given that bacteriophage are inert biological entities incapable of colonising in the absence of a sensitive host they could form a viable alternative that may have fewer safety implications and that could be delivered as a robust formulation.

## Background

The mammalian gastrointestinal tract is home to a complex and intimately associated microbial ecosystem (microbiome), comprised of bacterial (bacteriome), archaeal (archaeome), fungal (mycome) and viral (virome) components. The virome is mainly composed of bacteriophages (bacterial viruses collectively referred to as the phageome). It has been estimated that the human gut can contain as least as many bacteriophages as there are bacteria (10^14^) [1–3], making it perhaps the most densely populated ecological niche in nature. Co-evolution over millennia has selected those members of the microbiome that either cause no harm or confer a benefit to the host, these are commonly referred to as the commensals and mutualists of the gut [4–6]. A healthy microbiome is also associated with high bacterial strain level diversity [7] and bacteriophage may play an important role by restricting overgrowth of the most successful strains in ‘kill the winner’ dynamics. If phages have the potential to modulate the gut bacteriome, then in turn they could have an impact on host–microbe interactions and on host health [8]. Mammalian hosts have evolved to rely on microbial activities to assist digestion, provide vitamins, resist pathogens, and to regulate metabolism and the immune system [9] [10]. The intestinal microbiome is also a potent source of antigens and potentially harmful compounds [11]. Obtaining an appropriate microbiome at birth, and subsequently developing and maintaining it in the face of challenges such as antibiotic therapy, is an important determinant of health and wellbeing [12].

A multitude of studies have identified alterations in gut microbiomes in a wide variety of diseases, such as inflammatory bowel disease (IBD) including Crohn’s disease, activation of chronic human immunodeficiency virus (HIV) infection, atopy and a variety of psychological disorders, to name but a few [13–18]. Modernisation of society has impacted on microbiome diversity in many ways, including factors such as urbanization, global mobility, improved sanitation and hygiene, consumption of highly processed diets, non-familial child care, less contact with animals and medical therapies. It is likely that the ancestral relationship between a diverse gut microbiome and its human host has essentially been surpassed by the speed of invention and technology. A recent study that investigated the microbiome of the uncontacted Yanomami Amerindian people allows us a glimpse into the past. It appears that traditional lifestyles enabled the human gut to harbour a microbiome with high bacterial diversity and genetic function, something that has been compromised by modern lifestyles [19].

Antibiotics can be considered to be just one of these aspects of modernization and serves as an example of how essential medical therapies can lead to perturbations in the gut ecosystem, a situation exacerbated by misuse of antibiotics. Antibiotics have served as the initial line of defense against pathogenic bacteria for decades. The mass production of penicillin in the 1940s revolutionised modern medicine, ending a world where patients commonly died from bacterial endocarditis, bacterial meningitis, and pneumococcal pneumonia. Broad spectrum antibiotics have saved millions of lives but may leave the gut microbiome in an altered state in which the delicately constructed ecosystem of the gastrointestinal tract may be compromised.

Probiotics [20–22], prebiotics and other dietary interventions have been investigated [23] as tools to ameliorate the negative aspects of antibiotic use. The most radical therapy is faecal microbial transplantation (FMT), whereby the microbiome of an individual is supplemented by that of healthy donor [24]. In human therapy, FMTs are usually confined through practitioner guidelines to those suffering from severe, moderate or recurring *C. difficile* infections [25]. However, while case reports of FMT in patients with ulcerative colitis and Crohn’s disease are variable, some report that a certain percentage of patients achieve and can even maintain remission in some instances [26]. FMT has also been suggested as a potential therapy in irritable bowel syndrome, Type 2 diabetes mellitus, metabolic syndrome, obesity, fatty liver disease, multidrug resistant organism eradication, hepatic encephalopathy and paediatric allergy disorders [24].

With a standard FMT, the entire stool is homogenised and filtered through gauze and the resulting preparation is then infused into the gut. The entire microbiome is introduced into the recipient, including bacterial, archaeal, fungal and viral species. In this study we investigated if autochthonous virome transplantation (in which bacteriophage predominate) can impact on the bacteriomes and phageomes of mice following disruption with antibiotic treatment.

## Results

### Sequencing of the murine bacteriome

Two separate trials (designated as Study 1 and Study 2) were performed. In both trials 16S rRNA gene sequencing was used to determine the composition of the faecal bacteriome of pre!- and post-antibiotic treated mice (n=16). Half of the mice subsequently received an FVT (n=8) or a heat-treated FVT as a control (n=8). Virome transplantation was conducted using a bacteria free-virus faecal filtrate collected from the same mice prior to antibiotic treatment. In Study 1, a single gavage was performed after antibiotic washout, while in Study 2 a second gavage was also performed 4 days later (see experimental design in Fig S1). MiSeq (V3 kit - 300 bp paired end reads) sequencing resulted in an average of 16,805 ± 8,079 (Study 1) or 31,758 ± 64,605 (Study 2) paired end reads across the samples; and an average of 14,514 ± 7,095 (Study 1) or 29,350 ± 59,214 (Study 2) individual reads post-quality filtering (Supplementary Tables S1 and S2).

### Impact of antibiotics on the murine gut bacteriome

After a period of acclimatisation, mice were administered an antibiotic cocktail of penicillin and streptomycin in their drinking water for either four days (Study 1), or two days (Study 2). Following antibiotic treatment (CTAb) a significant change occurred in the murine bacteriome as compared to pre-treatment mice (samples at CT000 and CT00). These differences can be visualised via unweighted UniFrac PCoA (principal co-ordinate analysis; Fig 1 A), as well as via weighted UniFrac and Bray Curtis PCoA (Fig S2).

**Figure 1-.**
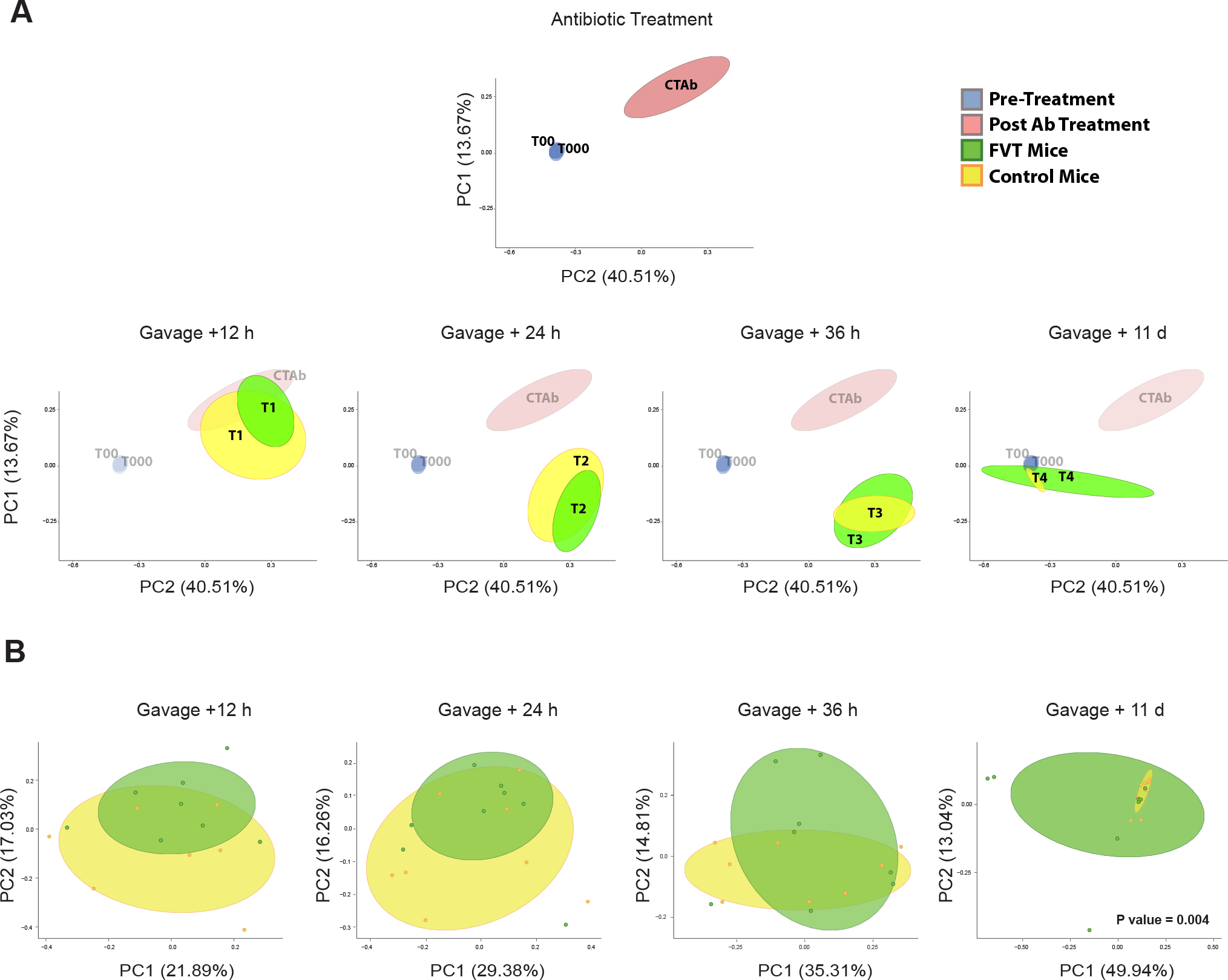
PCoA plots of unweighted UniFrac distances for Study 1 (A). Ellipses represent 70% confidence intervals. A UniFrac PERMANOVA test was performed with the Adonis function to determine the statistical differences between FVT and Control mice (Model formula = antibiotic treatment timepoint + Phage/control status), the resulting PCoA plots and P values are displayed (B).

Using the Shannon index as a measure of alpha diversity it is apparent that the bacterial diversity decreased after antibiotic treatment in both studies. In Study 1 (Fig S3A), after 4 days of antibiotic treatment (CTAb) there is a lower Shannon diversity than that observed for CTAb in Study 2 (Fig S3B), which is in consistent with a shorter two-day administration of antibiotics. However, when we examine bacterial cell numbers (log 16S copies/gram of faeces; Figure S4) there was a greater reduction in cell numbers in Study 2, when antibiotics were only administered for half the time (Study 1: log (16S copies/gram of faeces) mean 9.45; median 9.93; and Study 2: mean 7.80; median; 7.49). It must be noted however that in Study 2 the alpha diversity decreased further than that observed for the CTAb sample in the Control and FVT Mice in the T1 samples, and concomitantly bacterial numbers were elevated (Control mice T1: mean 8.56; median 8.76; and FVT Mice T1: mean 9.08; median 9.25) as was observed for Study 1-CTAb, where the lowest Shannon diversity of this study was also observed.

The increase in bacterial cell numbers may correlate with the outgrowth of certain species (Fig S5). For example, in Study 1 we observed an increase in *E. coli*/*Shigella* species in the antibiotic treated mice that continued to increase in the post gavage mice (CT1/VT1). After the shorter exposure to the antibiotic treatment in Study 2, this out-growth of *E.coli*/*Shigella* species was only observed in the faeces of the 10hr post-gavage (CT1/VT1) mice. In the CTAb mice, we did however observe an increase in “Other” species (those RSVs unclassified to genus level) and a decrease in *Alistipes*, *Bacteroides* and *Barnesiella* species. Classification of RSVs to a phylum level ascribes these changes to a decrease in Bacteroidetes and Firmicutes, and an increase in Actinobacteria, Proteobacteria, Cyanobacteria and unclassified others.

### The impact of bacteriophage on the antibiotic treated murine gut bacteriome

#### Study 1

Following antibiotic treatment mice were divided into two groups (n=8) and received a gavage of either autochthonous viable (FVT) or heat-treated bacteriophages (Control) by means of a faecal filtrate. In Study 1 the inferred PCoA plots of unweighted Unifrac distances (Fig 1A) we observe separation of the bacteriomes of Control and FVT mice into different clusters that both gradually return to a location similar to that of the pre-antibiotic bacteriome. Eleven days post-gavage there is a significant difference in the bacteriomes of FVT and Control groups as revealed by Adonis PERMANOVA analysis (P-value = 0.004). These differences were also observed using Bray Curtis distances but not using weighted Unifrac (Fig S2A, Table S3).

Differentially abundant taxa were also identified between the FVT and Control groups 11 days post-gavage (Fig 3A). Twelve RSVs were statistically distinct in their abundance (Table S4). Of these 12, six belong to the family Lachnospiraceae, while Ruminococcaceae accounted for one, as did Porphyromonadaceae and Deferribacteraceae. The final three were unclassified to the family level but are of the Order Clostridiales. Interestingly, all 12 RSVs not only differed in their abundance between CT4 vs VT4 mice but all are statistically different in their abundance compared to the pre-treatment mice (CT00). With 10 of these RSVs increased in Control mice (that did not receive viable phage), it implies that the FVT aided in restoring the murine bacteriome to its pre-treatment state. The RSVs increased in Control mice were of the predominantly of the family Lachnospiraceae, the remainder included the species *Mucispirillum schaedleri* and *Parabacteroides goldsteinii*. Only two RSVs were increased in mice receiving phage and these belong to the families Lachnospiraceae and Ruminococcaceae and include the species *Eubacterium siraeum*.

**Figure 2-.**
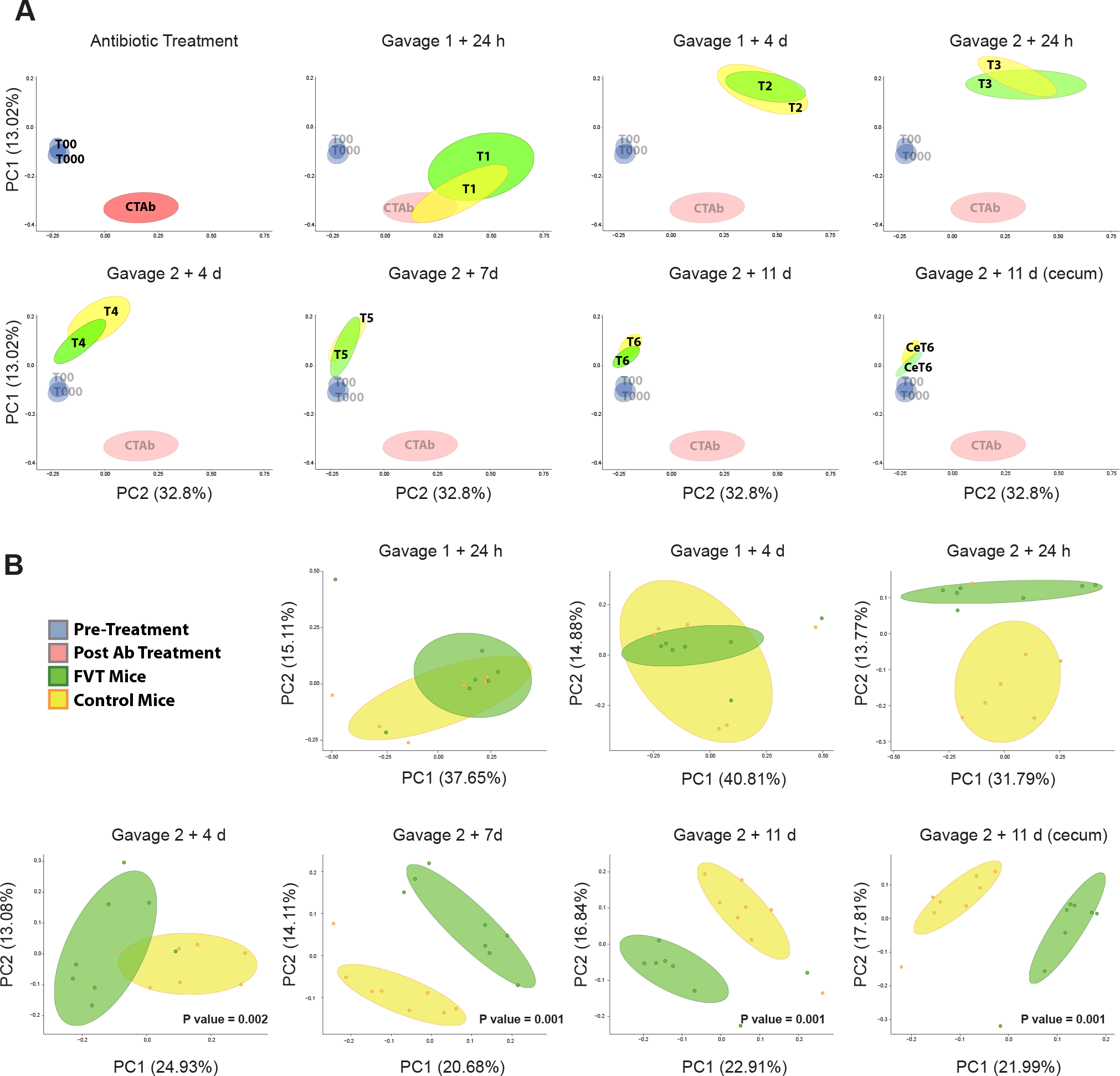
PCoA plots of unweighted UniFrac distances for Study 2 (A). Ellipses represent 70% confidence intervals. A UniFrac PERMANOVA test was performed with the Adonis function to determine the statistical differences between FVT and Control mice (Model formula = antibiotic treatment timepoint + Phage/control status), the resulting PCoA plots and P values are displayed (B).

**Figure 3-.**
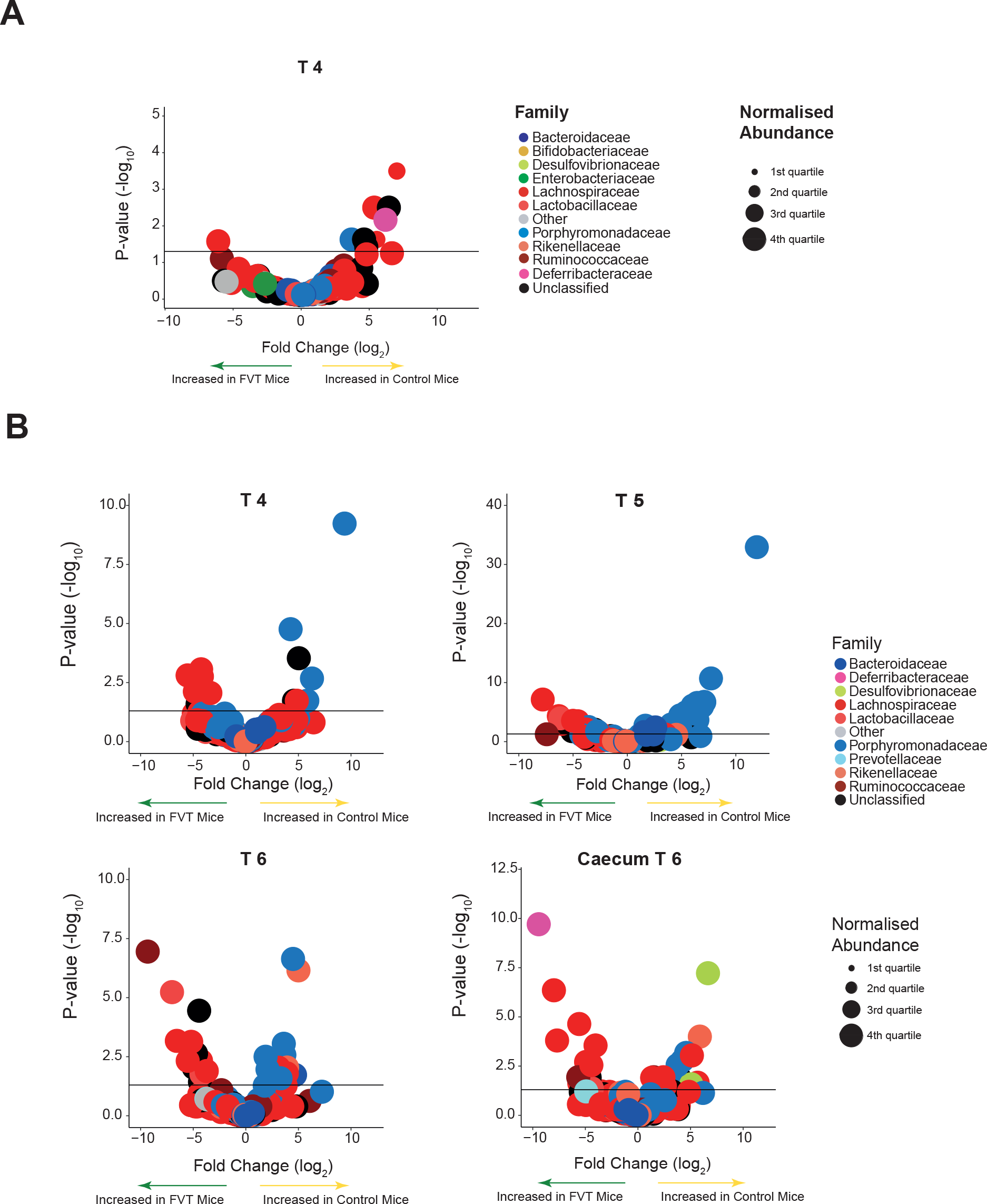
Volcano plots showing the results of DESeq2 which detects differentially abundant RSVs between the FVT and Control mice at each time point in Study 1 (A) and Study 2 (B). RSVs with an adjusted p-value <0.05 are positioned above the horizontal line. Normalised abundance is also represented. Families belonging to the phylum Firmicutes are indicated via varying shades of red, while families derived from Bacteroides are visualised in blue, additional colours are used to represent other families and phyla.

#### Study 2

Study 1 demonstrated that administration of a bacteriophage enriched gavage can impact the murine microbiota and so we performed a second more comprehensive study. Here a second antibiotic gavage was administered 4 days after the first and an increased number of samples (including caecum content) were analysed.

In Study 2, PCoA plots (Unweighted Unifrac, Bray Curtis and Weighted Unifrac (Fig 2A and S2B)) display clear clustering of RSVs defined by whether they received heat killed or viable phage. As in Study 1, over time the bacteriome of the mice gravitates from the post-antibiotic treatment state (CTAb) to the pre-treatment state (CT00 & CT000). We observed separation of the microbiota of mice into significantly different clusters from four days post-Gavage 2 (T4) to the end of the experiment (T6), where these differences were also revealed in the ceca (CeT6) (Adonis PERMANOVA: T4 P-value = 0.002, T5 P-value = 0.001, T6 P-value = 0.001, CeT6 P-value = 0.001) (Fig 2B). These differences were also observed using Bray Curtis distances but not in the case of weighted Unifrac (Fig S2B, Table S3).

Alpha diversity confirms a clear increase in diversity post-antibiotic treatment in both control and FVT receiving mice (Fig S3B). Several differentially abundant taxa were observed between CT4 and VT4 (13 RSVs), CT5 and PT5 (51 RSVs), CT6 and PT6 (37 RSVs) and CceT6 and PceT6 (25 RSVs) (Table S5). 26 RSVs were noted to be differentially abundant across two or more time points and five were highlighted to be differentially abundant at T6 in both faeces and caecum, revealing that these core taxa are more directly influenced by the presence or absence of administered bacteriophages. With respect to these 31 RSVs the taxa were almost equally distributed between Bacteroidetes (52%) and Firmicutes (48%). At the family level Porphyromonadaceae (36%) and Lachnospiraceae (32%) predominated. Lactobacillaceae accounted for a further 7% of RSVs, while Rikenellaceae and Ruminococcaceae accounted for 6% and 3%, respectively. The most commonly assigned genus was *Barnesiella* at 16%; however, 65% of RSVs could not be assigned to the genus level. Of the *Barnesiella* identified 75% were classified to be of the species *Barnesiella intestinihominis*.

A volcano plot was used to visualise the differential association of RSVs across the time points T4, T5, T6 and in the caecum at T6 (Fig 3B). Firmicutes family members (red hues) are more abundant in FVT animals across the final two time points and in the caecum, while Bacteroidetes family members (blue hues) are more abundant in Control mice. If we compare the faecal Firmicutes-Bacteroidetes ratio with pre-treatment (CT00) mice, we observe a significant decrease in this ratio in both sub-sets at T4. This ratio decreases further in Control mice at T6 while those that received bacteriophage return to that observed for pre-treatment mice at T6 (Kruskal-Wallis with Dunn’s multiple comparison post-test; CT00 vs CT4/VT4: P-value <0.05; CT00 vs CT6: P-value <0.01; CT00 vs VT6: P-value >0.05) (Fig 4).

**Figure 4 -.**
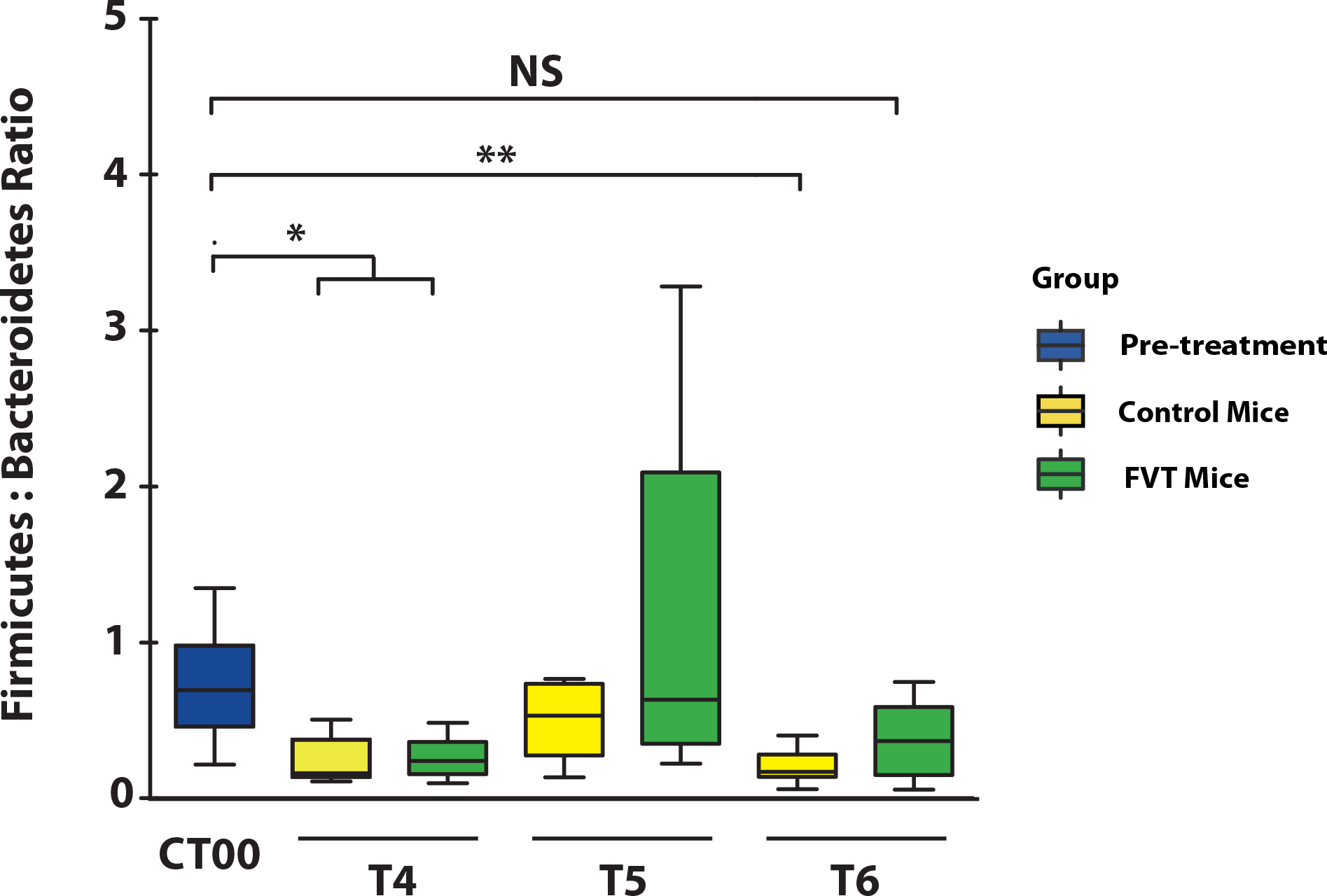
Bacteroidetes Firmicutes ratio reveals that the ratio is statistically higher in FVT mice as determined via Kruskal-Wallis with Dunn’s multiple comparison post-test; CT00 vs CT4/PT4: P-value <0.05; CT00 vs CT6:P-value <0.01; CT00 vs PT6: P-value >0.05)

Further examination revealed that certain taxa remain increased in FVT or Control mice over time (Table S5). For example, RSV_8 belonging to the family Porphyromonadaceae can be seen to fluctuate in abundance across the three faecal times points represented here, but is consistently increased in Control mice. All differentially abundant RSVs as determined by DESeq2 were compared to the abundance of these same RSVs in pre-treatment mice. Of the differentially abundant taxa at T4, 31% of the RSVs increased in FVT mice differed in their abundance as compared to pre-treatment (CT00) mice, while only 23% of differentially abundant RSVs in Control mice differed from pre-treatment mice. Over time these percentages decrease in bacteriophage-receiving FVT mice with only 10% at T5 and 5% at T6 of the RSVs differing in their abundance from CT00 mice, while in Control mice the opposite is observed with 12% at T5 and 54% at T6 (Supplementary table S5-RSVs that differ from CT00 mice are highlighted in blue).

### Murine viromes

In addition to examining the impact of bacteriophages on the murine bacteriome, we also performed a metagenomic study on DNA isolated from viruses extracted from the murine faecal samples. Technical limitations involving virome yield meant that the results represent the viromes of groups of mice rather than individual animals, in that faecal samples were pooled for pre-treatment (n=16), antibiotic treated (n=16), control (n=8) and FVT treated mice (n=8) at each time point (Fig S1). The virome data was analysed utilising an assembly based approach. Notably the viral sequences assembled here are not represented in current databases.

The virome that made up the FVT gavage can now be visualised as being present in the CT000 and CT00 samples (Fig 5). In both studies the virome composition differs between control and FVT mice in the post-gavage samples. Moreover, 11 days post antibiotic treatment those mice that received an FVT retain a notable abundance of the bacteriophages introduced in the gavage. For example, in Study 1 APC-pVirus10 is found in V-VT4 mice but is absent from the corresponding control V-CT4; in Study 2 APC-pVirus9 and APC-pVirus21 are found in abundance in V-VT6 mice where they are all but diminished in the control.

**Figure 5 -.**
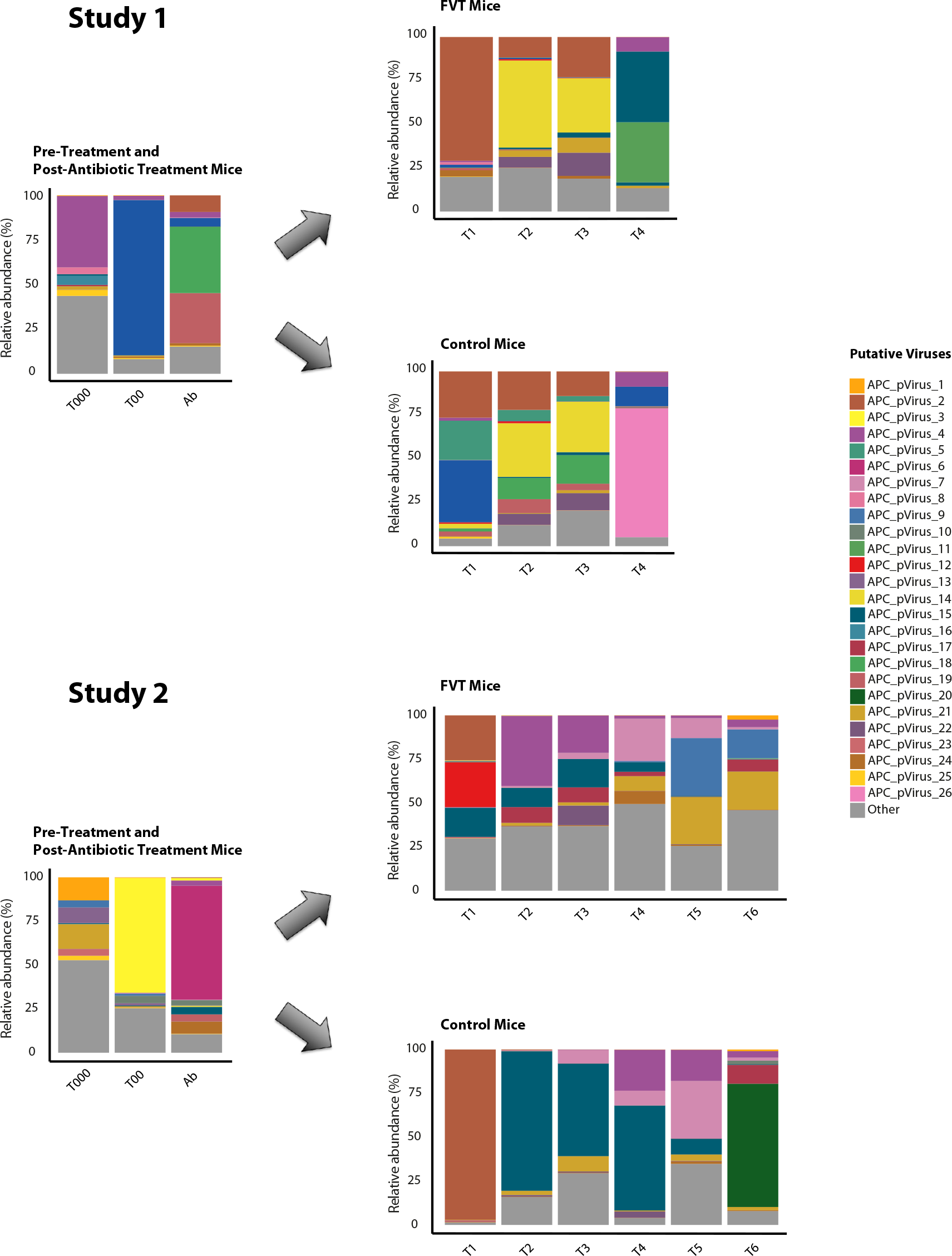
Metagenomic analysis on DNA isolated from viruses extracted from murine faecal samples in Study 1 (A) and Study 2 (B). Results reflect the viromes of groups of mice, where faecal samples were combined and represent the viral pool in pre-treatment (n=16), antibiotic treated (n=16), Control (n=8) and FVT mice (n=8) at each time point. Following sequencing contigs were assembled into putative viruses and the relative abundances are represented within this bar plot.

In conjunction with the outgrowth of *E. coli*/*Shigella* post-antibiotic treatment, we detected an increase in the abundance of APC-pVirus2. This is particularly noticeable in Control mice (V-CT1) that did not receive a viable FVT compared to those in receipt of FVT (V-VT1) (Fig 5 and Fig S5). Moreover, one putative bacteriophage APC-pVirus15 appears to be increased in abundance in the gut of Control mice compared to their FVT-receiving counterparts at corresponding time points. This is observed in both Study 1 and Study 2, suggesting the presence of particular bacterial hosts, common in these control animals, are facilitating replication of this bacteriophage.

Within the individual studies, we have reported that at specific time points differentially abundant bacterial taxa were detected. In Study 1, we observed that in the final time point (where such differentially abundant bacterial taxa were observed) that the phageomes of control and FVT mice differ not only in content but also in their corresponding relative abundances. A total of 152 bacteriophages were identified in the eight Control mice at this time point, while 163 are associated with FVT-receiving mice (Fig 5). With respect to viral abundances at this final time point, APC-pVirus11 and APC-pVirus15 constitute the majority of the viral load of Control mice, while APC-pVirus26 predominates in the gut of FVT mice. In Study 2, where 2 sequential gavages of viable FVT were administered, the variability in the profiles of the putative bacteriophages present and their relative abundances between control and FVT animals appears to differ even more. The richness of viral diversity is also elevated in FVT mice, especially in the final time points in tandem with the observation of differentially abundant bacterial taxa. This equates to a 20-30% increase in specific bacteriophages in these animals up to 11 days post gavage (see supplementary table S5). It must be noted however that these results are based on a pooled sample and so no statistical conclusions can be drawn.

## Discussion

The impact of externally added phages on GI tract bacteria has been studied for 100 years. d'Herelle in 1919 used a phage preparation to treat dysentery in what was almost certainly the first attempt to use bacteriophages therapeutically [42]. Since then there have been multiple studies revealing the impact of phage on a range of GI pathogens [for reviews see 8, 43–45]. Intestinal bacteriophages are likely to influence and shape the microbial ecology of the mammalian gut, similar to that observed in aquatic systems where it has been proposed that phages are responsible for 20–80% of total bacterial mortality and play a significant role in limiting bacterial populations [2, 46]. Bacteriophages have a profound impact on microbial population dynamics and are one of the driving forces shaping the functionality and diversity of the human intestinal microbiome [reviewed in 2].

Here we report on how the administration of an autochthonous FVT had a significant impact on the bacteriome of mice following antibiotic-induced disruption. The bacteriome and phageome of FVT-receiving animals differed in the abundance and/or diversity of species. While the benefits of faecal transplantation have been well documented, the impact of bacteriophage within this microbial community has yet to be fully investigated. The transfer and long-term colonisation of bacteriophages species during standard FMT has been documented [47], with temperate phage members of *Siphoviridae* found to be transferred between human donors and recipients with significantly greater efficiency than other bacteriophage groups [48]. Bacteriophages may well be important elements of FMT that can influence and shape gut ecology. One study has in fact indicated that a sterile filtered faecal transplant (FFT), where cellular microbes were removed, produced longitudinal changes in the bacterial and viral community structures of faecal samples in five human recipients suffering from *Clostridium difficile* infections, all of whom recovered following treatment [49]. The potential role of bacteriophage in the recovery of the microbiota from the ravaging effects of this infection was acknowledged, along with potential roles for bacterial components and metabolites.

In the two studies described here, we disrupted the murine microbiome with antibiotics prior to administration of autochthonous bacteriophage. These antibiotics have a broad spectrum of activity. Streptomycin is an aminoglycoside inhibitor of protein synthesis that binds primarily to 16S ribosomal RNA and inhibits both Gram-positive and Gram-negative bacteria. Penicillin is a β-lactam inhibits cell wall biosynthesis and targets Gram-positive species. These antibiotics have also been shown to exhibit synergistic activity [50, 51]. Previous studies have shown that treatment of mice with streptomycin increases susceptibility to oral infection with *Salmonella enterica* serovar Typhimurium [52]. After antibiotic treatment we observed an outgrowth in *E. coli/Shigella* species that coincided with low α-diversity and elevated bacterial cell numbers. Such outgrowth could be linked with antibiotic insensitivity of these species allowing them to dominate their more sensitive relatives. This has been described for *E. coli* in a number of studies [53, 54]. The occurrence of these elevated levels of *E. coli* varies across the two studies and perhaps can be attributed to deviations in the length of antibiotic treatment pre-sampling.

Following the administration of a single FVT gavage in Study 1, we witnessed an alteration in the faecal bacteriome in PCoA plots and via the presence of differentially abundant taxa at the final time-point. This impact re-shaping of the gut bacteriome was also observed, and to a greater extent, during Study 2 where the bacteriophage dose was doubled by means of sequential gavages four days apart. In Study 2 differentially abundant taxa were observed across the final three time-points and also in the ceca. Samples derived from the murine caecum have a bacteriome composition distinct from that of the faecal samples [55]. Therefore, differences observed here between phage treated and Control mice indicates alterations in the bacteriome at multiple GI sites and infers that the impact of bacteriophage on the murine gut is widespread.

Each gram of human faeces contains approximately 10^11^ bacterial cells, ~ 10^11^ bacteriophages, ~10^7^ colonocytes, ~10^8^ archaea, ~10^8^ viruses, ~10^6^ fungi, protists and metabolites [56–58]. The protocols used to process the murine faeces used to prepare the FVT gavage in this study would have removed all biological species apart from bacteriophages and viruses (and those metabolites or microbial structures are also present in the control gavage). Heat inactivation of bacteriophage and the use of nucleases ensures that the Control mice would have only received small-molecule metabolites, bacterial components or antimicrobial compounds of bacterial origin (e.g. bacteriocins) that contribute to the normal intestinal microenvironment. Thus, we can deduce that the alterations to microbiome (both bacteriome and virome) of test mice is due to the acquisition of viable bacteriophages.

Of the differentially abundant RSVs identified, a subset were annotated as the species *Barnesiella intestinihominis*, which appeared on multiple occasions to be present in greater abundance in Control mice (Study 2 - T4, T5, T6 and caecum T6). This Gram-negative commensal bacterium has been classified as an oncomicrobiotic for its immunomodulatory properties [59]. Loss of this species in FVT-receiving mice suggests that they obtained a bacteriophage that could utilise *B. intestinihominis* as a host, thus diminishing the population of this species. This is described as the “kill the winner” model, but what also may be the case is the “kill the relative” model, whereby the induction or presence of prophage/lysogenic bacteriophages can trigger an epidemic among susceptible bacterial competitors [57]. Several other models that describe how bacteriophage can shift the dynamic of bacterial populations also exist. One model of interest and relevance to this study is the “community shuffling model”, which involves the induction of prophage and stimulation of the host immune system by environmental factors [57] such as sub-inhibitory levels of antibiotics, such as quinolones or beta-lactams (as was utilised here albeit at inhibitory levels) from bacterial species such as *E. coli* [60], *Clostridium difficile* [61], *E. faecalis* [62] and *Staphylococcus aureus* [63]. Should the presence of antibiotics within the current studies have encouraged such events, it should have occurred prior to administration of exogenous bacteriophages and so would have been present in both control and test animals. These prophages could however have contributed to the bacteriophage profile present in CTAb mice and to the initial ecology to which the administered FVT was introduced.

Given that the FVT treated mice had a distinct bacteriome, we were able to identify a number of differentially abundant bacterial taxa and these could be visualised on volcano plots. Via colour coding of family members, it became apparent that there was a clear shift in favour of Firmicutes family members in those that received bacteriophages. This may indicate that these mice did not have or receive bacteriophages that target these family members, in particular the Lachnospiraceae. Alternatively, there may be other population dynamics models at play such as those mentioned previously. Conversely, in Control mice such bacteriophage either flourished or the ecological conditions of the gut were so that members of the Bacteroidetes family predominated. Firmicutes-Bacteroidetes ratios confirmed that this phenomenon was not confined to differentially abundant taxa but was statistically valid across all reads.

When we examine how the presence/absence of differentially abundant bacterial taxa compare to these same RSVs in pre-treatment mice after acclimatisation (CT00), we observed that the bacteriome of FVT mice more closely resembles that of CT00 mice than that of the Controls. This indicates that autochthonous bacteriophages have impacted and shaped the bacteriome, restoring it to a pre-treatment state. By killing bacteria, bacteriophages implement population fluctuations in their bacterial hosts, they significantly influence global biochemical cycles, and are also considered to be crucial in driving microbial species diversity due to the fact that they are species-specific [64, 65].

Within these studies we pooled faecal matter from CT000 and CT00 mice (pre-antibiotic treatment) and created a sterile autochthonous faecal virome for transplantation. We used metagenomic DNA sequencing and assembly of the resulting contigs to determine the bacteriophage content. In both studies, FVT mice appeared to retain the bacteriophage they received up to 11 days post gavage, long after the transit time of several hours noted for bacteriophage through the GI tract [66]. While we cannot draw any statistical conclusions since we only had a single autochthonous combined faecal sample for each group, our observations mirror those observed by Ott et al., 2016 [49] in humans that received an FFT, with the recipient virome altering substantially and retaining features of the FVT sample.

Gut bacteriophages are so numerous and diverse that sequence databases contain only a small fraction of the total diversity [67–69], and so we used an assembly approach to identify these bacteriophages rather than relying on annotation via alignment to public databases. Of the bacteriophages identified here, none are currently present on public viral databases, we propose that this assembly approach opens the door to discovery of unknown bacteriophages and viruses whose impact and influence in the microbiome could go unnoticed simply due to the paucity of current viral databases.

## Conclusions

To our knowledge this is the first time that an autochthonous FVT has been shown to re-model and restore the mammalian gut following a period of microbial disruption. While others such as Ott et al, 2016 have suggested the role of bacteriophage in human FFT (fecal filtrate transplant; [49]), this study with the inclusion of a control treatment has validated a role for bacteriophage in microbiome population dynamics. There is potential to take advantage of this observation, especially with the increasing trend toward performing FMTs. Until now these have largely focussed on transferring living bacteria and spores; but bacteriophage are non-living proteinaceous entities that could form a robust, inexpensive alternative that could be delivered as a freeze dried formulation.

## Methods

### Mouse models and experimental design

#### Study1

16 BALB/c mice were obtained from Harlan Laboratories UK Ltd. and were housed within the Biological Services Unit, University College Cork (UCC). Mice were received at 7-8 weeks of age and allowed to acclimatise for 5 days on a standard rodent diet. During this time faecal samples were collected, frozen (−80°C) and used, in part, to prepare FVT phage-rich material which will be used for oral gavage. This required obtaining 2-3g of faecal pellets which were resuspended in a solution of 10ml filter sterilised SM buffer (50 mM Tris-HCl; 100 mM NaCl; 8.5 mM MgSO_4_; pH 7.5) and 2ml 1M NaHCO_3_ (to help deacidify the stomach). After vortexing, the solution is centrifuged at 4700rpm for 20mins. The supernatant was then filter sterilised (0.45μM filter) and divided into 2 separate containers. One to be administered to the test (FVT) group and contains active bacteriophages/viruses, while the second was heated at 95°C for 15 mins, followed by DNase (Ambion) treatment at 37°C for 1 hr (according to manufacturer’s guidelines) as described by Reyes et al [27], and thus contained inactivated bacteriophages to be administered to the control group. After acclimatisation, all 16 mice were administered antibiotic treatment (penicillin 1000U/ml and streptomycin 3g/L) in their drinking water for 4 days (and faecal samples were collected) followed by 1 day of antibiotic wash-out, where standard drinking water was administrated *ad libitum*. Following this, the mice were divided into 2 groups (n=8), faecal samples were collected and the FVT group were gavaged with 0.2ml of the FVT material (as prepared above) while the control group received inactivated bacteriophages. Faecal samples were then collected 10hr, 24hr, 34hr and 11 days afterward.

#### Study 2

Study 2 was performed as that described for Study 1 with the following deviations. 16 BALB/c mice were received at 8-10 weeks of age and allowed to acclimatise for 3 days. After acclimatisation, all 16 mice were administered antibiotic treatment (penicillin 1000U/ml and streptomycin 3g/L) in their drinking water for 2 days followed by 1 day of antibiotic wash-out. Following this, the mice were divided into 2 groups (n=8), faecal samples were collected and the FVT group were gavaged with 0.2ml of active bacteriophages (as prepared above) while the control group received inactivated bacteriophages (“Gavage 1”). Faecal samples were then collected 1 and 4 days afterward. Forthwith a second gavage (“Gavage 2”) was subsequently administered in a similar manner as described for Gavage 1. Again faecal samples were collected from both groups post-gavage at 1, 4, 7 and 14 days. Post-mortem, caecum content was also collected.

### DNA extractions and library preparation for MiSeq

Faecal samples were all frozen immediately after collection at times indicated in Fig S1 and then used to extract bacterial DNA for 16S rRNA analysis of the bacteriome (~2-5 faecal pellets were collected from each mouse at each time point and used track alterations in the bacteriome of each individual mouse). The QIAamp DNA Stool Mini Kit (Qiagen, Hilden, Germany) was used according to manufacturer’s guidelines to extract bacterial DNA but was modified to include a bead-beating step. 16S ribosomal DNA hypervariable regions V3 and V4 were amplified via PCR using a high fidelity polymerase (Phusion; Thermo Fisher Scientific) and the primers V3F - 5’-CCTACGGGNGGCWGCAG-3’ and V4R – 5’-GACTACHVGGGTATCTAATCC-3’ [28] with the addition of the appropriate Illumina Nextera XT overhang adapter sequences (Illumina, San Diego, CA, USA). Following purification using a magnetic bead capture kit (Ampure; Agencourt), the amplicon libraries underwent a second PCR reaction to attach dual indices and Illumina sequencing adapters using the Nextera XT index kit (Illumina, San Diego, CA, USA). Following purification (as described above) the dsDNA libraries were quantified using a Qubit^®^ Fluorometer (Thermo Fisher Scientific) were then pooled in equimolar concentrations. Ready to load libraries were sequenced on an Illumina MiSeq (Illumina, San Diego, California) using V3 sequencing kit (300 bp paired end reads) at GATC Biotech AG, Germany.

### Analysis of 16S sequencing data

The quality of the raw reads was visualized with FastQC v0.11.3. T Forward and reverse reads in were merged using FLASH [29] Primers were removed from the merged reads using the fastx_truncate command of USEARCH [30]. The trimmed reads were then demultiplexed with a Phred quality score of 19 using the split_libraries_fastq.py script of the QIIME package [31]. The demultiplexed sequences in FASTA format were then de-replicated using the derep_fulllength command of USEARCH and sequences below a minimum length cutoff (400nt) were removed. Singletons were removed using the minsize option of the sortbysize command of USEARCH. The resulting sequences were then clustered into OTUs using the cluster_otus command and then chimera filtered using both the de novo and reference based chimera filtering implemented in USEARCH with the ChimeraSlayer gold database v20110519 [30] The reads were then aligned back to the OTU’s using the usearch_global command of USEARCH to generate a count table which was input into R for statistical analyses. Taxonomy was assigned to the sequences using mothur v1.38 [32] against the RDP database version 11.4, as well as classified with SPINGO to species level [33]. Only ribosomal sequence variants (RSVs) with a domain classification of Bacteria or Archaea were kept for further analysis. A phylogenetic tree of the RSV sequences rooted on the midpoint was generated with FastTree [34]. Alpha diversity and Beta diversity were generated using PhyloSeq v1.16.2, which also was used for a principle coordinates analysis as implemented in Ape v3.5. Differential abundance analysis was carried out with DESeq2 v1.12.4[35]. All visualisation in R was performed with ggplot2 v2.2.1. Permutational multivariate analysis of variance (PERMANOVA) was performed in R Vegan package with the adonis function (Model formula = antibiotic treatment timepoint + Phage/control status) with 9999 permutations.

### Real-time qPCR

A LightCycler^®^ 480 apparatus (Roche), associated with LightCycler^®^ 480 Software (version 1.5; Roche), was used for the real-time PCR. Each reaction contained a 5μl of a 1 in 10 dilution of genomic DNA and was carried out in quadruplicate in a volume of 15 μl in a 384-well LightCycler^®^ 480 PCR plates (Roche), sealed with LightCycler^®^ 480 sealing foil (Roche). Amplification reactions were carried out with Phusion 2X master mix (Thermo Fisher Scientific) using run conditions, primers (0.5pmol each per reaction) and probe (0.1pmol per reaction) as described by Furet et el [36]. Quantitation was done by using standard curves made from known concentrations of linearized plasmid DNA containing the 16S amplicon. Wells containing nuclease free water were included as negative controls. Statistical analysis was performed using Graphpad Prism 5, whereby a One-Way ANOVA followed by Tukey test determined statistical significance; ***P value<0.001, ** P value<0.01, *P value<0.05.

### Virome DNA extraction and library preparation for MiSeq

DNA corresponding to the viromes of each group of mice was purified from faecal samples, with approximately 1 pellet per mouse included in the extraction. Samples were taken at the time points indicated in Fig S1. Faecal samples were homogenised in 10ml SM buffer followed by centrifugation twice at 5,000g at 10°C for 10mins and filtration through a 0.45μm syringe filter to remove particulates and bacterial cells. NaCl (0.5M final concentration; Sigma) and 10% w/v polyethylene glycol (PEG-8000; Sigma) were added to the resulting filtrate and incubated at 4°C overnight. Following centrifugation at 5,000g at 4°C for 20mins, the pellet was resuspended in 400μl SM buffer. An equal volume of chloroform (Fisher) was added and following 30 sec of vortexing the sample was centrifuged at 2,500g for 5 mins at RT. The aqueous top layer is retained and it was subjected to RNase I (10U final concentration; Ambion) and DNase (20U final concentration; TURBO DNA-free™ Kit, Invitrogen) treatment in accordance with the manufacturer’s guidelines. To isolate DNA, virus like particles were incubated with 20 μL of 10% SDS and 2 μL of proteinase K (Sigma, 20 mg/mL) for 20 min at 56 °C, prior to lysis by the addition of 100 μL of Phage Lysis Buffer (4.5 M guanidine thiocyanate; 45 mM sodium citrate; 250 mM sodium lauroyl sarcosinate; 562.5 mM β-mercaptoethanol; pH 7.0) with incubation at 65°C for 10 min. Viral DNA was purified by two treatments with an equal volume of phenol:chloroform:isoamyl alcohol (25:24:1) and passing the resulting purified DNA through a QIAGEN Blood and Tissue Purification Kit and eluting samples in 50 μL of AE Buffer. In study 1 the viral DNA was used directly for Nextera XT library preparation (Illumina) as described by the manufacturer. In Study 2 the DNA concentrations were equalised prior to amplification using an Illustra GenomiPhi V2 kit (GE Healthcare). Amplifications of purified viral DNA was performed in triplicate on all samples as described by the manufacturer. Subsequently, an equal volume of each amplification and an equal volume of the original viral DNA purification were pooled together and used for paired-end Nextera XT library preparation. All samples were sequenced on an Illumina MiSeq at GATC in Germany.

### Analysis of virome sequencing data

The quality of the raw reads was visualized with FastQC v0.11.3. Nextera adapters were removed with cutadapt v1.9.1 [37] followed by read trimming and filtering with Trimmomatic v0.36 [38] to ensure a minimum length of 60, maximum length of 250, and a sliding window that cuts a read once the average quality in a window size of 4 follows below a Phred score of 30. Reads were then assembled with the metaSPAdes assembler [39]. In order for a contig to be included in the final analysis it must have been at least 5kb in length, then either detected as viral by Virsorter [40] in the virome decontamation mode, or had a significant BLAST hit (95% identity over 95% of the length) to a genome in RefSeq Virus, or had no significant BLAST hits (any alignment length with an e-value greater than 1e-10) against nt. This allowed us to include known viruses, putative viruses predicted by virsorter and completely novel viral sequences not yet included in any database. The quality filtered reads were then aligned to this contig set using bowtie2 v2.1.0 [41] using the end to end alignment mode. A count table was generated with samtools v0.1.19 which was then imported into R v3.3.0 where the relative abundance of contigs (labelled as putative viruses) was plotted using ggplot2 v2.2.1.

## List of Abbreviations

FVT: faecal virome transplantation
FFT: fecal filtrate transplant
FMT: faecal microbial transplantation
PCoA: principle co-ordinate analysis
IBD: inflammatory bowel disease
HIV: human immunodeficiency virus

## Declarations

### Ethics approval and consent to participate

All procedures involving animals were approved by the Irish Department of Health and Children and performed by an individual authorised to do so under licence number B100/3729.

### Consent for publication

All authors consent to this publication.

### Availability of data and material

All sequence data used in the analyses were deposited in the Sequence read Archive (SRA) (http://www.ncbi.nlm.nih.gov/sra) under BioProject PRJNA385256 for the 16S rRNA sequence data and BioProject PRJNA385134 for the virome sequence data. Sample IDs, meta data and corresponding accession numbers are summarized in supplementary table S7. All raw count tables, 16S taxonomic assignments and R code used for the analysis are available at https://figshare.com/s/eb1666b10037656b987d

### Authors’ contributions

Conceived and supervised the study: CH, and RR. Designed the experiments: MD, CH, and RR. Performed the experiments: PC, MD and LD. Analyzed the data: LD, FR, MD, VV, and AM. Wrote the paper: LD, FR, RR and CH.

## Acknowledgements

Not applicable

---

**Figure S1-.**
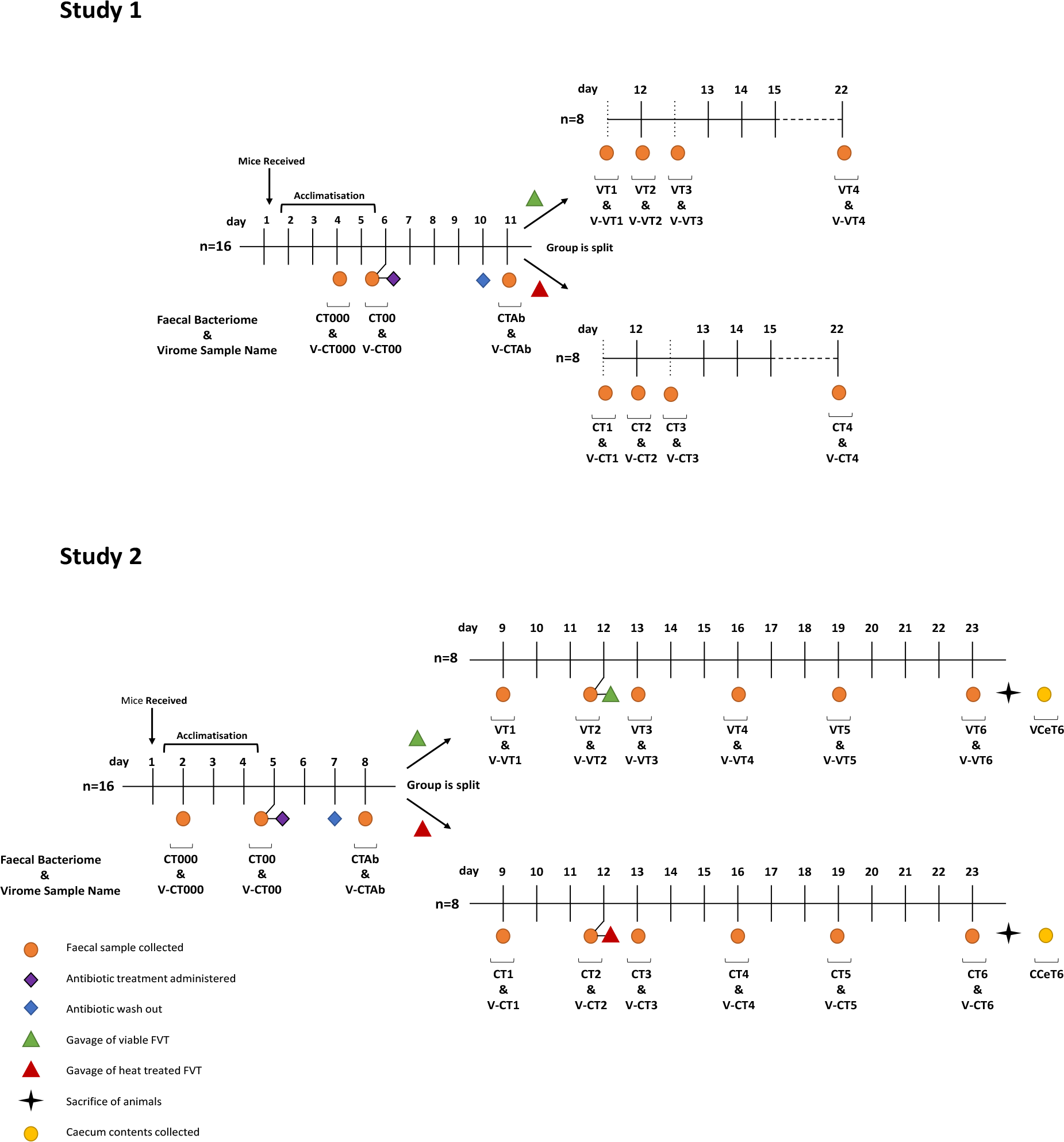
Experimental design of Study 1 (A) and Study 2 (B). BALB/c mice (n = 16) were, after acclimatisation, administered antibiotic treatment. After a period of antibiotic wash-out the group was split into two (n=8) and the mice were gavaged with an FVT, a sterile virome faecal filtrate (either viable or heat killed) that had been isolated from frozen faecal samples obtained from the mice during acclimatisation. In Study 2 (B) a second gavage was administered 4 days after the first, dotted vertical lines representing time points within a day. Each group of mice was individually caged. Each solid vertical line represents a day. Time points selected for sampling the faecal microbiota of each mouse in each treatment group are represented as circles and labelled. Samples were subjected to 16S rRNA sequencing and viral metagenomic sequencing.

**Figure S2-.**
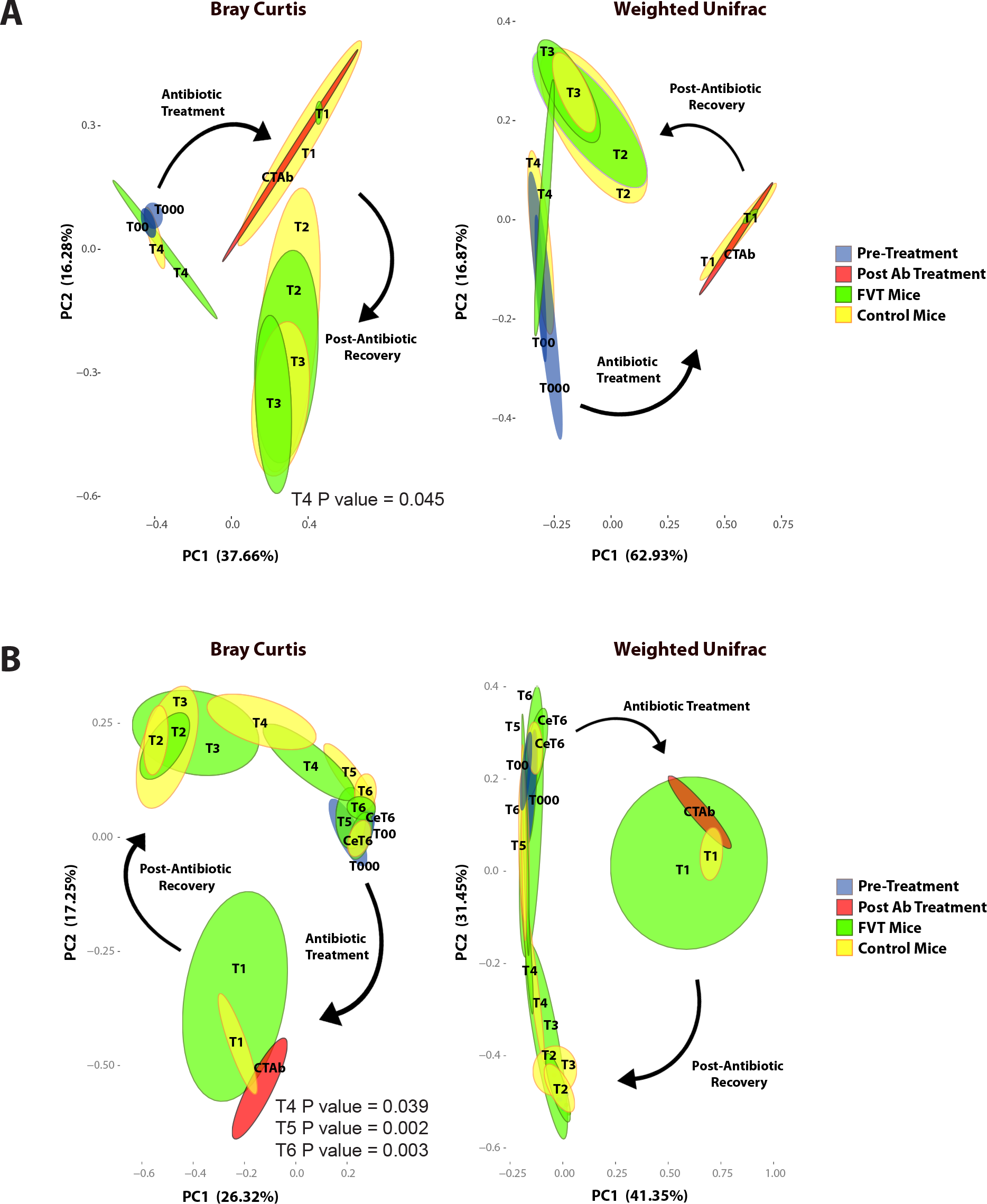
PCoA plots compiled using Bray Curtis and Weighted Unifrac for Study 1 (A) and Study 2 (B). Statistically significant P values following UniFrac PERMANOVA analysis performed with the Adonis function to determine the statistical differences between FVT and Control mice have been inserted.

**Figure S3-.**
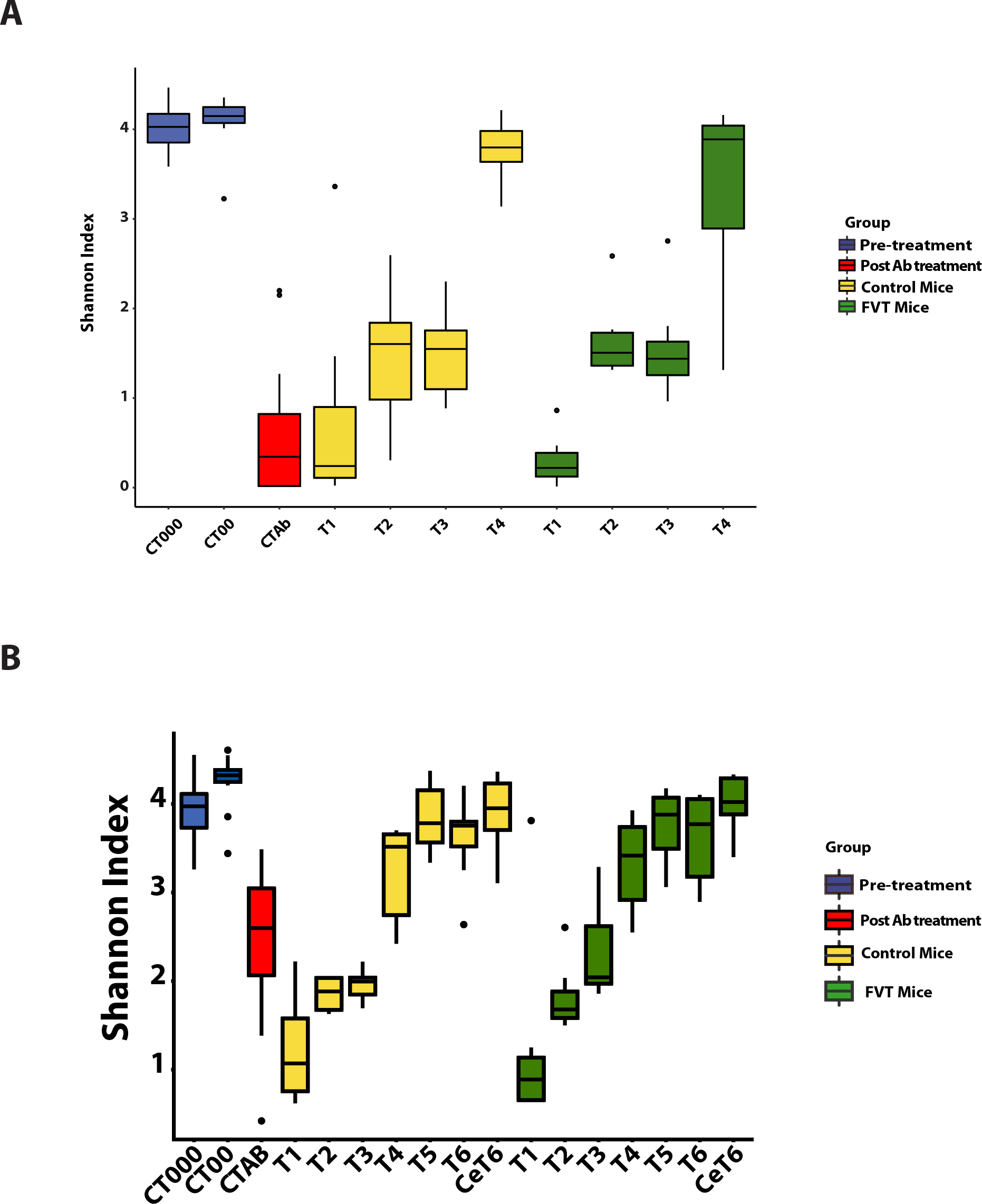
Shannon diversity index was used to display the bacteriome alpha diversity over time for Study 1 (A) and Study 2 (B). No statistical differences were observed in alpha diversity between FVT and Control mice at corresponding time points.

**Figure S4-.**
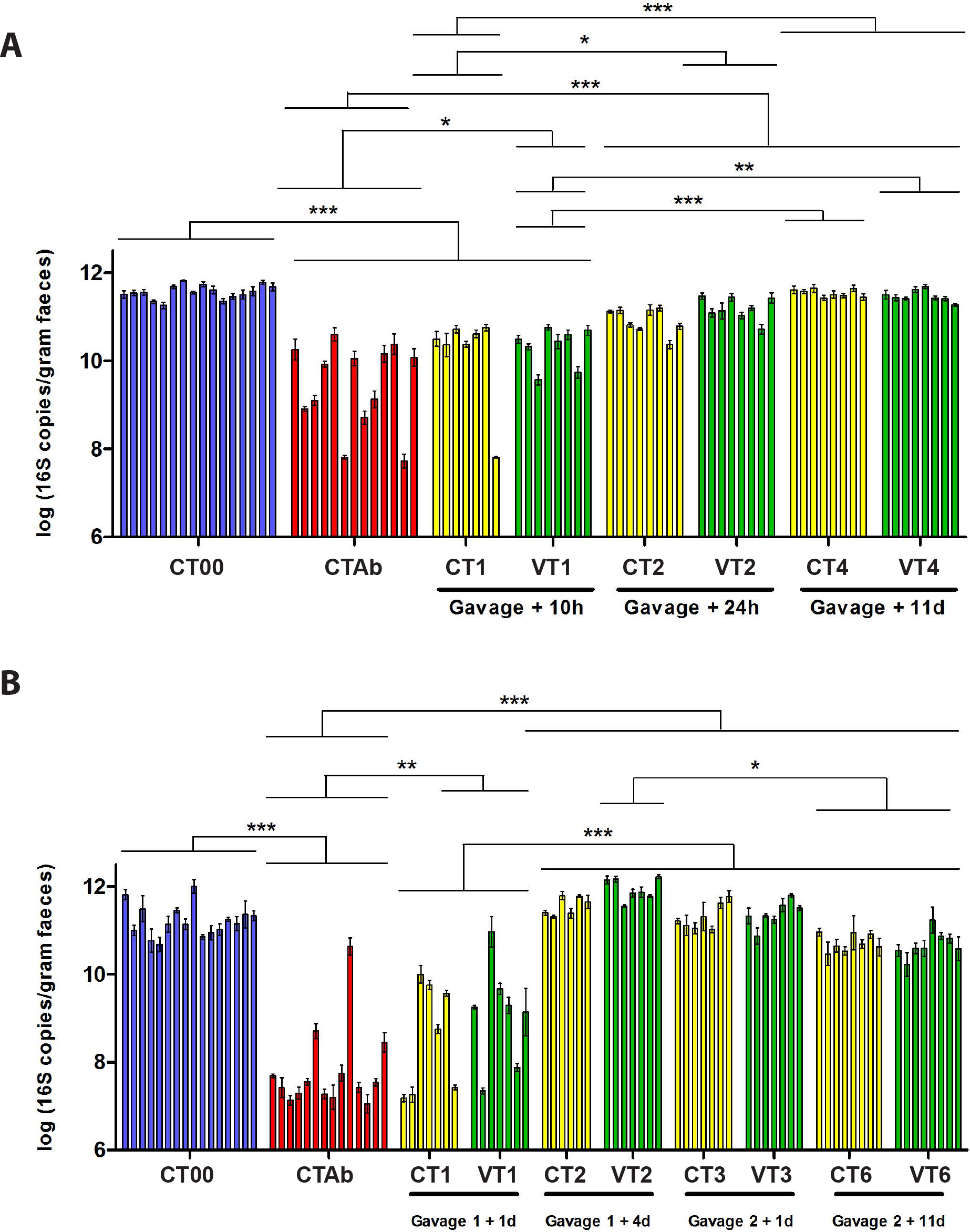
qPCR was used to determine the approximate bacterial cell numbers present per gram of faeces in Study 1 (A) and Study 2 (B). Bacterial cell numbers were seen to drop dramatically following antibiotic treatment in both studies. A One-Way ANOVA followed by Tukey test determined statistical significance; ***P value<0.001, ** P value<0.01, *P value<0.05.

**Figure S5-.**
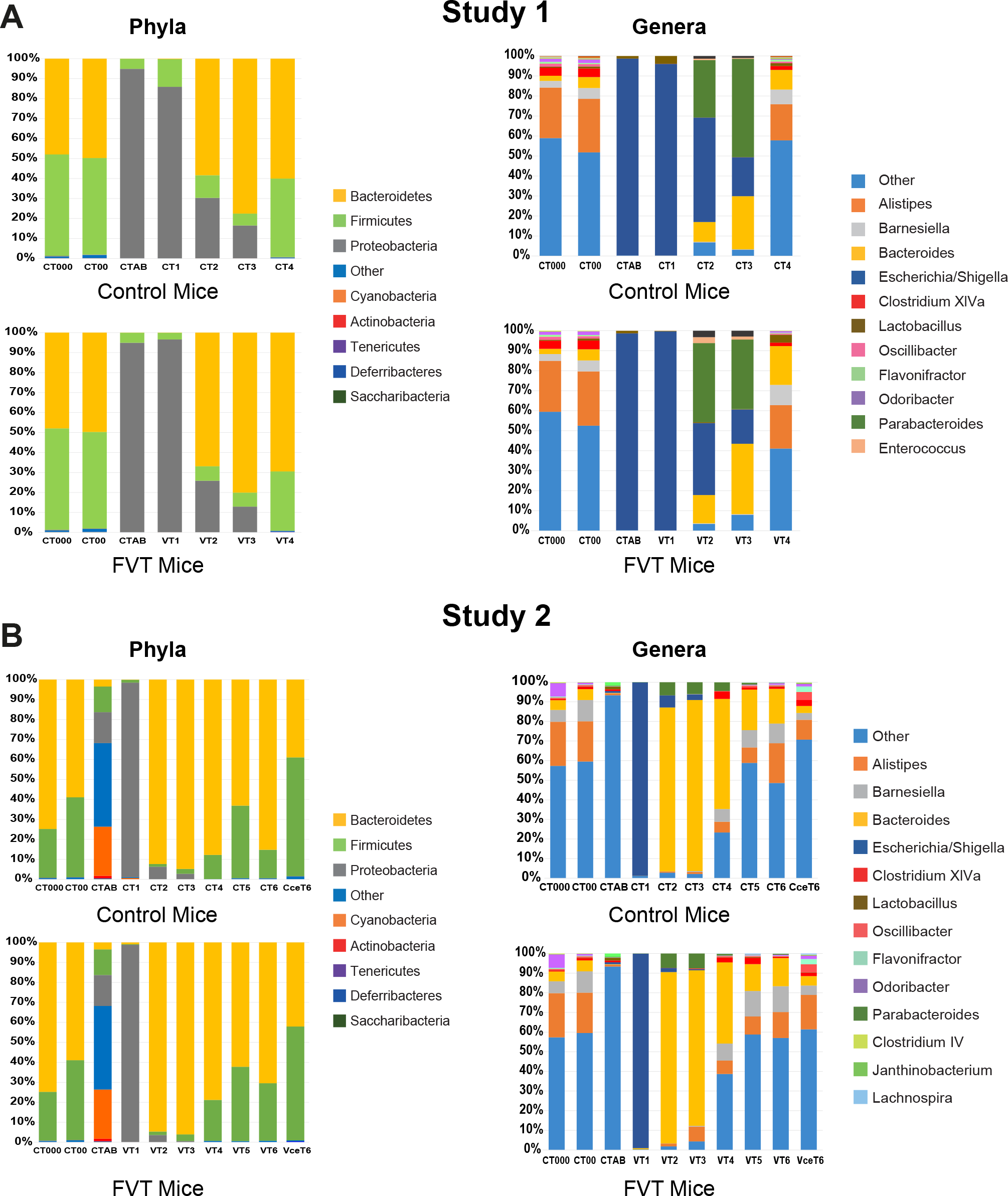
The abundance of different taxa visualised here at the genus level for Study 1 (A) and Study 2 (B) in mice pre-treatment (CT000 and CT00), post-antibiotic treatment (CTAB) and over time post-gavage with either viable bacteriophage (Study 1: VT1-VT4; Study 2: VT1-VceT6), or heat treated non-viable bacteriophage (Study 1: CT1-4; Study 2: CT1-CceT6).

